# Comparative analysis of L-fucose utilization and its impact on growth and survival of *Campylobacter* isolates

**DOI:** 10.1101/2021.09.09.459711

**Authors:** Pjotr S. Middendorf, Wilma F. Jacobs-Reitsma, Aldert L. Zomer, Heidy M. W. den Besten, Tjakko Abee

## Abstract

*Campylobacter jejuni* and *Campylobacter coli* were previously considered asaccharolytic, but are now known to possess specific saccharides metabolization pathways, including L-fucose. To investigate the influence of the L-fucose utilization cluster on *Campylobacter* growth, survival and metabolism, we performed comparative genotyping and phenotyping of the *C. jejuni* reference isolate NCTC11168 (human isolate), *C. jejuni* Ca1352 (chicken meat isolate), *C. jejuni* Ca2426 (sheep isolate), and *C. coli* Ca0121 (pig manure isolate), that all possess the L-fucose utilization cluster.

All isolates showed enhanced survival and prolonged spiral cell morphology in aging cultures up to day seven in L-fucose-enriched MEMα medium (MEMαF) compared to MEMα. HPLC analysis indicated L-fucose utilization linked to acetate, lactate, pyruvate and succinate production, confirming the activation of the L-fucose pathway in these isolates. Highest consumption of L-fucose by *C. coli* Ca0121, is conceivably linked to its enhanced growth performance up to day 7, reaching 9.3 log CFU/ml compared to approximately 8.3 log CFU/ml for the *C. jejuni* isolates. Genetic analysis of their respective L-fucose clusters revealed several differences, including a 1 bp deletion in the *Cj0489* gene of *C. jejuni* NCTC11168, causing a frameshift in this isolate resulting in two separate genes, *Cj0489* and *Cj0490*, while no apparent phenotype could be linked to the presumed frameshift in the NCTC11168 isolate. Additionally, we found that the L-fucose cluster of *C. coli* Ca0121 was most distant from *C. jejuni* NCTC11168, but confirmation of links to L-fucose metabolism associated phenotypic traits in *C. coli* versus *C. jejuni* isolates requires further studies.

**Importance:** *Campylobacter* is the leading cause of gastroenteritis in humans worldwide, with increasing incidence and prevalence in recent years. The most prevalent species are *Campylobacter jejuni* and *C. coli* with 83% and 10% of all *Campylobacter* cases, respectively. Previously it was found that the majority of *Campylobacter* isolates are able to metabolize L-fucose (fuc+ isolates), a sugar that is widely present in the human gut. Putative roles for L-fucose in fuc+ *C. jejuni* isolates were found in growth, biofilm formation and virulence. Despite this, relatively little is known about L-fucose metabolism and the impact on growth and survival in fuc+ *Campylobacter* isolates. The results from our comparative genotyping and phenotyping study demonstrate that L-fucose, in both *C. jejuni* and *C. coli* fuc+ isolates, is involved in enhanced survival, prolonged spiral cell morphology and changes in the general metabolism. Possible links between phenotypes and differences in respective L-fucose gene clusters are discussed.

## Introduction

*Campylobacter* is a leading cause of gastroenteritis in humans worldwide (1, 2). The incidence and prevalence of campylobacteriosis has increased over the past few years in both developed and developing countries (3–6). Most common *Campylobacter* species causing gastroenteritis are *Campylobacter jejuni* and *Campylobacter coli*, causing 83% and 10% of all human cases, respectively (3,7). Studies reported that *C. jejuni* can infect and cause diarrhea with a relatively low infection dose, generally causing symptoms like gastroenteritis with acute watery or bloody diarrhea, fever and abdominal pain (4,8–10). Post-infection complications include the severe Guillain-Barré syndrome and Miller-Fisher syndrome (11). Most infections are related to incidents that have been reported due to consumption of meat products predominantly poultry, direct contact with animals and via environmental waters (12–20).

Interestingly, *Campylobacter* is generally recognized as being susceptible to a wide variety of environmental stresses (21–25). However, outside the animal and human gastro-intestinal tract, *Campylobacter* is able to survive and has therefore a certain degree of environmental robustness which is needed to endure environmental transmission. Nutrient availability and acquisition support *Campylobacter* transmission between different animal hosts and the human host (26,27). Several studies dedicated to the characterization of substrate utilization in *Campylobacter* spp. showed an important role for citric acid cycle intermediates, amino acids and peptides in supporting growth (28–30). Preferred amino acids used by *C. jejuni* include serine, aspartate, asparagine, and glutamate (31–33). *Campylobacter* was previously thought to be asaccharolytic, lacking most key enzymes to metabolize sugars. However, more recently, evidence was provided that selected *C. coli* and *C. jejuni* isolates can metabolize glucose and/or L-fucose (34–39). A systematic search for Entner–Doudoroff (ED) pathway genes encoding glucose utilization enzymes in a wide range of *C. coli* and *C. jejuni* isolates from clinical, environmental and animal sources, and in the *C. jejuni/coli* PubMLST database, revealed that 1.7% of the more than 6,000 available genomes encoded a complete ED pathway involved in glucose metabolism (39). Based on additional phenotyping, it was concluded that some glucose-utilizing *C. coli* and *C. jejuni* isolates exhibit specific fitness advantages, including stationary-phase survival and biofilm production, highlighting key physiological benefits of this pathway in addition to energy conservation (30,39).

Notably, comparative WGS analysis revealed that approximately 65% of the sequenced *C. jejuni* isolates and 73% of the sequenced *C. coli* isolates possess the L-fucose utilization cluster (designated *Cj0480c* – *Cj0490* in *C. jejuni* NCTC11168), so called fuc+ isolates. The L-fucose utilization cluster is regulated by *Cj0480c* and contains 2 predicted transporters encoded by *Cj0484* and *Cj0486*/*FucP*, of which only *Cj0486* was functionally linked to L-fucose metabolism (37). Uptake of L-fucose into the cell leads to its conversion into α-L-fucose, which is further transformed into β-L-fucose via a predicted mutarotase (*Cj0488*), and subsequently converted into L-fuconolactone and L-fuconic acid via a short chain dehydrogenase (*Cj0485/FucX*) and a amidohydrolase (*Cj0487*) (35,37). The next step in the pathway is the transformation of L-fuconic acid into 2-oxo-3-deoxy-L-fuconic acid via an altronate hydrolase/dehydratase (Cj0482/*Cj0483*) and its subsequent hydrolyzation into pyruvic acid and lactaldehyde by a dihydrodipicolinate synthase (*Cj0481/DapA*) with the latter compound converted to lactic acid by aldehyde dehydrogenase (*Cj0489*/*Cj0490*) (35,37).

Putative roles for L-fucose in fuc+ *C. jejuni* isolates in growth, biofilm formation and virulence have been reported, however to date, no data is available about the effect of L-fucose on *C. coli* (34,36,37). The human gastro-intestinal tract is a fucose-rich environment with fucose incorporated in glycan structures found on epithelial cells (40,41). These fucosylated glycans can be hydrolyzed by fucosidases produced by a range of bacterial inhabitants of the gut such as *Bacteroides* spp.. *C. jejuni* does not possess any obvious fucosidase homologs, however, *C. jejuni* can forage on L-fucose released by *Bacteroides vulgatus* in co-cultures with porcine mucus as a substrate (35). Furthermore, fucosylated glycans are not only commonly used as carbon source but can also serve as adhesion sites or receptors for pathogens like *Helicobacter pylori* and *C. jejuni* (42). Previous research in *Campylobacter* has shown that fucose monomers and fucosylated glycans serve as chemoattractant for *C. jejuni*, supporting adherence to epithelial cell surfaces containing such glycans (43,44).

Despite the potential role of L-fucose in the intestinal ecology and infection efficacy of *C. jejuni*, relatively little is known about the activation of L-fucose metabolism and the impact on stationary phase survival in fuc+ *Campylobacter* isolates. Interestingly, in human disease *C. coli* is less prevalent in comparison with *C. jejuni*, however, the L-fucose utilization cluster is more common among *C. coli* isolates (34). A comparative analysis of the efficacy and metabolite formation following activation of L-fucose utilization clusters in *C. jejuni* and *C. coli* isolates, combined with impact on growth and survival, has not been reported up to now.

In this study we investigated the impact of L-fucose on growth and long-term culturability of three fuc*+ C. jejuni* isolates and one fuc+ *C. coli* isolate, and we correlated this to the utilization of L-fucose and genetic features of the L-fucose genomic cluster in the tested isolates.

## 2. Methods

### 2.1 Bacterial isolates and culture preparation

The following *Campylobacter* isolates were used during this study*: C. jejuni* NCTC11168 (reference isolate) isolated from human feces, *C. jejuni* Ca1352 isolated from chicken meat, *C. jejuni* Ca2426 isolated from sheep manure and *C. coli* Ca0121 isolated from pig manure. *Campylobacter* stock cultures were prepared using BactoTM Heart Infusion broth (Becton, Dickinson and Company, Vianen, the Netherlands) and were grown for 24 h at 41.5 °C in microaerobic conditions (5% O_2_, 10% CO_2_, 85% N_2_) which was created using an Anoxomat WS9000 (Mart Microbiology, Drachten, the Netherlands). Glycerol stocks were prepared using 30% glycerol and 70% overnight culture and they were stored at −80 °C.

Routinely, prior to growth experiments, the Campylobacter freezer stocks were streaked on Columbia Agar Base (CAB (Oxoid, Landsmeer, the Netherlands)) plates supplemented with 5% (v/v) lysed horse blood (BioTrading Benelux B.V. Mijdrecht, the Netherlands) and 0.5% bacteriological agar No.1 (Oxoid) for optimal recovery. The plates were incubated in anaerobic jars for 24 h at 41.5 °C in microaerobic conditions. Colonies were routinely selected and grown overnight in 10 mL GibcoTM MEMα medium (Thermo Fisher Scientific, Bleiswijk, the Netherlands) (see Table. S1 for medium details) supplemented with 20 μM FeSO4 (Merck Schiphol-Rijk, the Netherlands), 10 mL MH2 broth (Merck) or 10 mL MH3 broth (Oxoid). A second overnight culture was made by diluting the Campylobacter suspension 1:100 in 10 mL fresh MEMα, MH2 or MH3 medium. The suspension was incubated at 41.5 °C for 24 h in microaerobic conditions to obtain standardized working cultures for use in further experiments.

### 2.2 L-fucose growth experiments

Prior to growth experiments, MEMα medium was supplemented with 21.0 mM L-fucose (MEMαF medium) and filter-sterilized using 0.2-μm pore sized filters. Infusing bottles were closed using a rubber stopper and aluminum cap and next they were sterilized. Sterilized infusion bottles were filled with 45 mL filter-sterilized MEMα medium or MEMαF medium by using a syringe. Filled infusion bottles were stored at 4 °C until further use.

Working cultures were decimally diluted in MEMα medium to a cell concentration of approximately 10^5^ CFU/mL. A final dilution step was done by adding 5 mL into the infusion bottles filled with 45 mL MEMα medium or MEMαF medium, resulting in a starting cell concentration of 10^4^ CFU/mL. Incubation of the inoculated infusion bottles was done at 37 °C. Daily, approximately 4 mL sample was taken from each infusion bottle, starting at day 0. After each sampling point, the head space of infusion bottles was flushed for 2 min with microaerobic gas (5% O_2_, 10% CO_2_, 85% N_2_) using a home-made gas flushing device using syringes to puncture the rubber stopper.

The samples were used to determine the bacterial concentration and for microscopic analysis. The remainder of each sample was stored at −20 °C for high pressure liquid chromatography (HPLC) analyses. Bacterial concentrations were determined by decimally diluting 1 mL of sample in peptone physiological salt solution (PPS, Tritium Microbiologie, Eindhoven, the Netherlands), followed by surface plating on CAB plates. CAB plates were incubated in jars for 48 h at 41.5 °C in microaerobic conditions. Colonies were counted and expressed in log_10_ CFU/mL. Each sample was microscopically analyzed using an Olympus BX 41 microscope (lens Ach 100x/1.25, Olympus Nederland, Leiderdorp, the Netherlands) and pictures were captured using CellSens Imaging software (Olympus Corporation). Three biologically independent reproductions were performed per condition, i.e. MEMα medium, and MEMαF medium, on different days.

### 2.3 High pressure liquid chromatography for organic acids

Samples obtained from the growth experiments were centrifugated at 13,000 *g* at 4°C for 5 min. Pellets were removed and the supernatant was treated for protein decontamination with Carrez A (K_4_FeCN)_6_.3H_2_0, Merck) and B (Z_n_O_4_·7H_2_0, Merck). After centrifugation, the supernatant was added to HPLC vials. Quantitative analyses were done using standards with pre-made concentrations for L-fucose, acetate, alpha-ketoglutarate, succinate, pyruvate and lactate. The HPLC was performed on an Ultimate 3000 HPLC (Dionex, Sunnyvile, USA) equipped with an RI-101 refractive index detector (Shodex, Kawasaki, Japan), an autosampler and an ion-exclusion Aminex HPX – 87H column (7.8 × 300 mm) with a guard column (Bio-Rad, Hercules, CA). As mobile phase, 5 mM H_2_SO_4_ (Merck) was used at a flow rate of 0.6 mL/min. Column temperature was kept at 40 °C. For each run, the injection volume was 10 μL and the run time 30 min. Chromeleon software (Thermo Fisher Scientific, Waltham, USA) was used for quantification of compound concentrations.

### 2.4 High pressure liquid chromatography for amino acids

Samples obtained from the growth experimented were used in aliquots of 40 μL. These aliquots were kept on ice and were diluted with 50 μL of 0.1 M HCl (containing 250 μM Norvalin as internal standard, Merck). The samples were deproteinized by addition of 10 μL of cold 5-sulphosalicilic acid (SSA, Merck) (300 mg/ml) and centrifuged at 13,000 *g* at 4°C for 10 min. In order to obtain an optimal pH for derivatization (pH between 8.2 to 10.0), approximately 60 to 150 μL of 4N NaOH was added to 5 mL of the AccQ•Tag™ Ultra borate buffer (Borate/NaOH buffer, Waters, Milford, USA). For derivatization 60 μL of Borate/NaOH was added to a total recovery vial. Twenty μL of the supernatant obtained after deproteinization of the plasma was added and mixed. To each of the vials 20 μL of AccQ Tag Ultra derivatization reagent (Waters) dissolved in acetonitrile was added and mixed for 10 s. Each vial was immediately capped. The vials were then heated for 10 min at 55 °C. The vials were stored at −20°C prior to HPLC analysis. Quantitative analyses were done using standards with pre-made concentrations for, histidine, asparagine, serine, glutamine, arginine, glycine, aspartic acid, glutamic acid, threonine, alanine, proline, cysteine, lysine, tyrosine, methionine, and valine. HPLC was performed on an Ultimate 3000 HPLC (Dionex) equipped with an RI-101 refractive index detector (Shodex), an autosampler and an ion-exclusion Aminex HPX – 87H column (7.8 × 300 mm) with a guard column (Bio-Rad). As mobile phase, eluants A and B (Waters) was used at a flow rate of 0.7 mL/min. Column temperature was kept at 55 °C. For each run, the injection volume was 1 μL and the run time 17 min. Chromeleon software (Thermo Fisher Scientific) was used for the determination of compound concentrations. Baseline separation was obtained for all amino acids except glutamine and arginine.

### 2.5 Genomic analyses

The sequence of NCTC11168 was obtained from the public collection of genbank with accession number AL111168. The sequences of the assembled genomes of *C. coli* Ca0121, *C. jejuni* Ca1352 and *C. jejuni* Ca2426 were obtained via the Netherlands Food and Consumer Product Safety Authority (NVWA) and are described in (45).

Genome alignments were performed using the online Benchling software (https://www.benchling.com). Each gene was translated into an amino acid sequence which was used for further alignments. Protein interactions were analysed with STRING (https://string-db.org/). Alignments were visualized by using the T-Coffee (46) or clinker (https://github.com/gamcil/clinker). Box shade figures were generated using BOX-SHADE 3.21 (https://embnet.vital-it.ch/software/BOX_form.html) using the RTF_old output format.

### 2.6 Statistical analyses

Differences in log_10_-counts observed for growth in MEMα medium and growth in MEMαF medium were statistically tested using a two-tailed Student’s *t*-test. P values ≤ 0.05 were considered as significant difference.

## Results

### Diversity in growth performance of *Campylobacter* in the absence and presence of L-fucose

We investigated growth and survival of different *C. jejuni* host isolates and one *C. coli* host isolate; namely *C. jejuni* NCTC11168 (human stool isolate), *C. jejuni* Ca1352 (chicken meat isolate), *C. jejuni* Ca2426 (sheep manure isolate) and *C. coli* Ca0121 (pig manure isolate), in MEMα medium and MEMαF medium up to 7 days. At day 1 and 2, no difference was observed between growth in MEMαF medium and MEMα medium for the tested isolates and at day 2 cell concentrations reached were 8.4, 8.2, 8.3 and 8.6 log_10_ CFU/mL, for *C. jejuni* NCTC11168, Ca1352, Ca2426 and *C. coli* Ca0121, respectively (Fig. 1A-D). However, from day 3 onwards, cell viability decreased in the absence of L-fucose and significantly lower cell counts were observed from day 3 until day 7, at which final cell counts reached 1.0, 4.4, 5.0 and 5.7 log_10_ CFU/mL in MEMα medium for isolates *C. jejuni* NCTC11168, Ca2426, Ca1352 and *C. coli* Ca0121, respectively. In MEMαF, cell concentrations did not decrease up to day 4 or 5, after which lower or stable cell concentrations were observed for isolate *C. jejuni* NCTC11168, Ca2426 and Ca1352 (Fig. 1A-C). Notably, *C. coli* Ca0121 showed continuation of growth after day 2, reaching 9.3 log_10_ CFU/ml at day 7 (Fig. 1D). Linking cell counts to morphology using microscopic images revealed that in MEMα medium, coccoid cells were commonly observed, increasingly over time after day 2, while in MEMαF medium, higher quantities of spiral-shaped cells were observed for all tested isolates (Fig. S1). In line with the observed increase in plate counts, isolate *C. coli* Ca0121 showed almost no increase in the amount of coccoid cells overtime when grown in MEMαF medium (Fig. S1).

**Figure 1.**
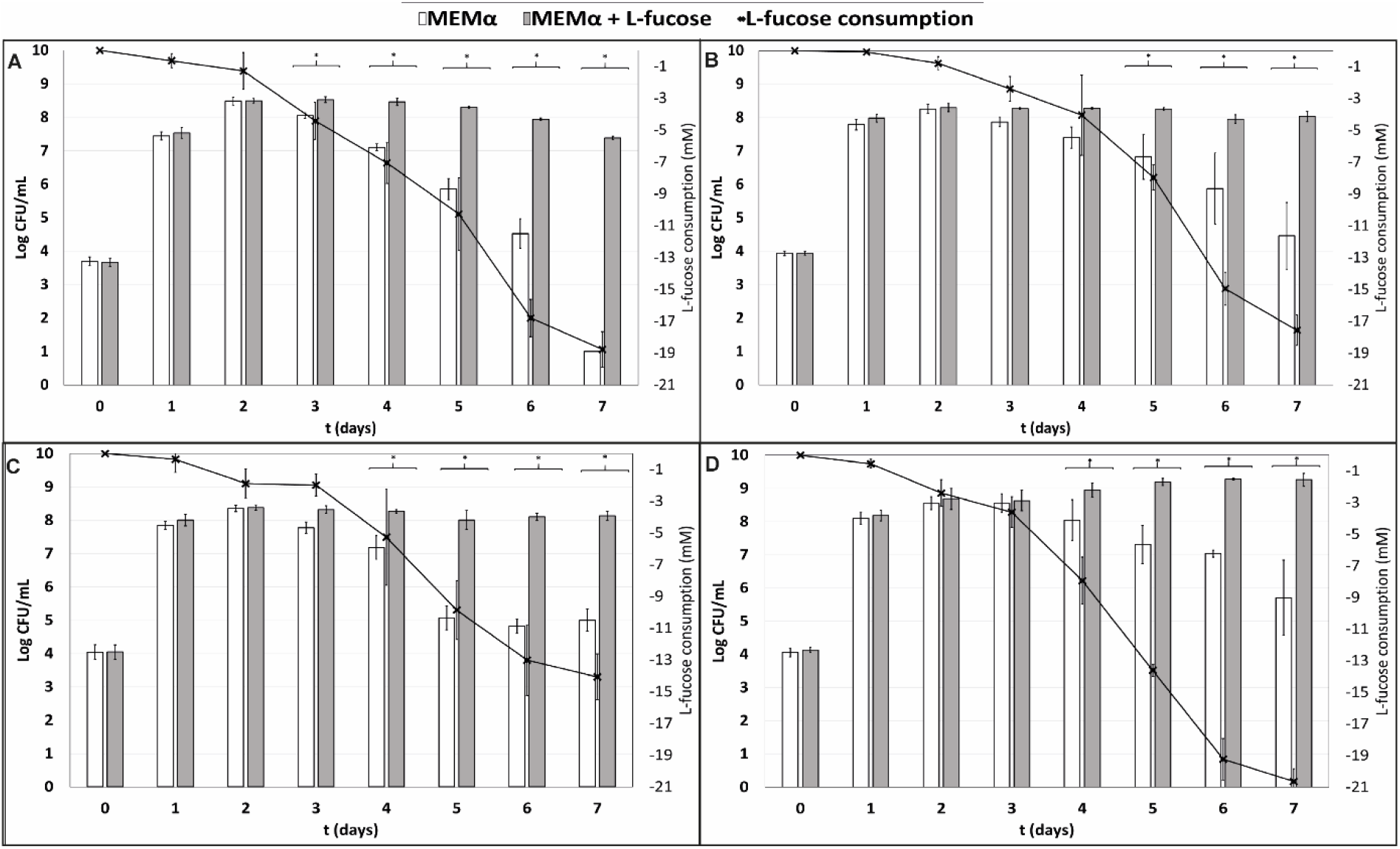
Quantification of planktonic growth for **A)** isolate *C. jejuni* NCTC11168 (human stool isolate), **B)** *C. jejuni* Ca1352 (chicken meat isolate) **C)** *C. jejuni* Ca2426 (sheep manure isolate) and **D)** *C. coli* Ca0121 (pig manure isolate), in MEMα medium (white bars) and MEMαF medium (grey bars). The black line shows consumption of L-fucose over time. Each value represents the average of 3 biologically independent replicates, and error bars show the standard deviation. The asterisks indicate significant (*P*<0.05) differences in cell counts after incubation in MEMα medium and MEMαF medium.

HPLC analyses confirmed absence of L-fucose in MEMα medium and enabled quantitative analysis of L-fucose consumption by the tested isolates in MEMαF medium. The L-fucose concentration decreased after day 1 for all four isolates. Interestingly, the *C. coli* isolate showed the highest and fasted L-fucose consumption over time, which was in line with the robust growth performance of this isolate.

### L-fucose consumption and metabolite concentrations in *C. jejuni* and *C. coli* isolates

We determined the concentrations of lactate, pyruvate, acetate, succinate and alpha-ketoglutarate during growth, as these have been associated with Campylobacter metabolism (27,35,47). Lactate production was not detected in non-supplemented MEMα medium after growth (Fig. 2). In MEMαF medium, up to 1.1 to 1.75 mM lactate was produced till day 5 and remained rather stable afterwards (Fig. 2). Starting concentrations of 0.25 mM of pyruvate were found to be depleted at day 2 in MEMα medium, whereas in MEMαF medium pyruvate concentrations remained at these low levels in *C. jejuni* NCTC11168 and *C. coli* Ca0121, and increased up to 0.6 mM and 1.3 mM in *C. jejuni* Ca1352 and Ca2426, respectively (Fig. 2). Also changes in acetate concentrations were observed. In MEMα medium low concentrations of acetate (<1 mM) were initially produced by all tested isolates and consumed later on (Fig. 2). In MEMαF medium, higher amounts of acetate were produced and concentrations increased until day 7 in isolates *C. jejuni* NCTC11168, Ca1352 and Ca2426 acetate reaching 4.4 mM, 4.1 mM and 2.9 mM, respectively. Notably, *C. coli* Ca0121 produced highest levels of acetate at day 5 (6.3 mM), and the lower level at day 7 pointed to acetate consumption. We observed a slight increase in succinate concentrations up to day 3, reaching 0.25 mM with *C. jejuni* isolates NCTC11168, Ca1352 and Ca2426, followed by consumption of succinate in both media (Fig. 2). Notably, *C. coli* Ca0121 produced highest levels of succinate up to day 3 in MEMα and up to day 5 in MEMαF, reaching 0.6 mM and 0.8 mM, respectively, after which succinate levels decreased in both media. Concentrations of alpha-ketoglutarate (6.5 mM) remained the same in all tested conditions (data not shown).

**Figure 2.**
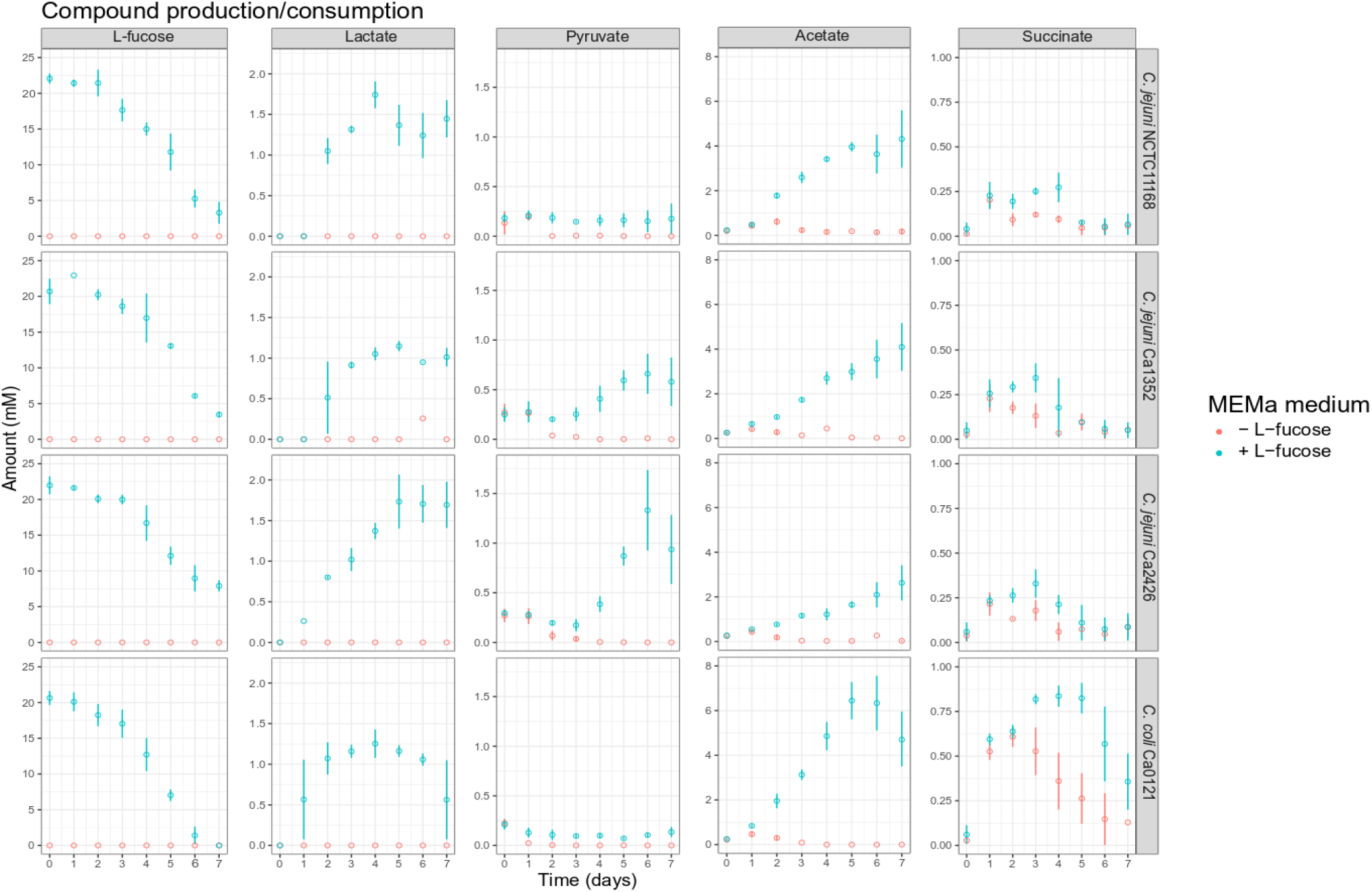
Compound production and consumption of selected *Campylobacter* isolates in MEMαF medium (blue dots) and MEMα medium (red dots) after a 7-day incubation period. Columns correspond to different compounds and each row shows results for a different isolate. Each value represents the average of three biologically independent replicates, and error bars show the standard deviation.

### Isolate diversity in *C. jejuni* amino acid metabolism

We investigated amino acid metabolism in all four tested isolates in MEMα medium and MEMαF medium and all 20 essential amino acids were quantified during the 7-day period (Fig 3. and Fig. S3). A strong preference was observed in all tested isolates for the amino acids serine and aspartic acid, with serine already being fully depleted from the medium after one day of incubation in MEMα medium for isolates Ca1352, Ca2426 and *C. coli* Ca0121 (Fig. 3). Isolates NCTC11168, Ca1352 and Ca2426 showed a transient peak of aspartic acid at day 2, that was rapidly depleted the following days. Glutamic acid was fully depleted after 3 days of incubation in all tested isolates without and with L-fucose (Fig. 3), while asparagine decreased in the same period but remained present at low levels (0.1-0.15 mM). Proline was also fully depleted in a time-dependent manner in all tested isolates and media, with *C. coli* Ca0121 showing a delayed consumption. Notably, a strong depletion of the sulphur-containing metabolite cystine in MEMαF medium with all *C. jejuni* isolates was observed in the first 2 or 3 days, whereas no significant consumption was observed in MEMα medium (Fig. 3).

**Figure 3.**
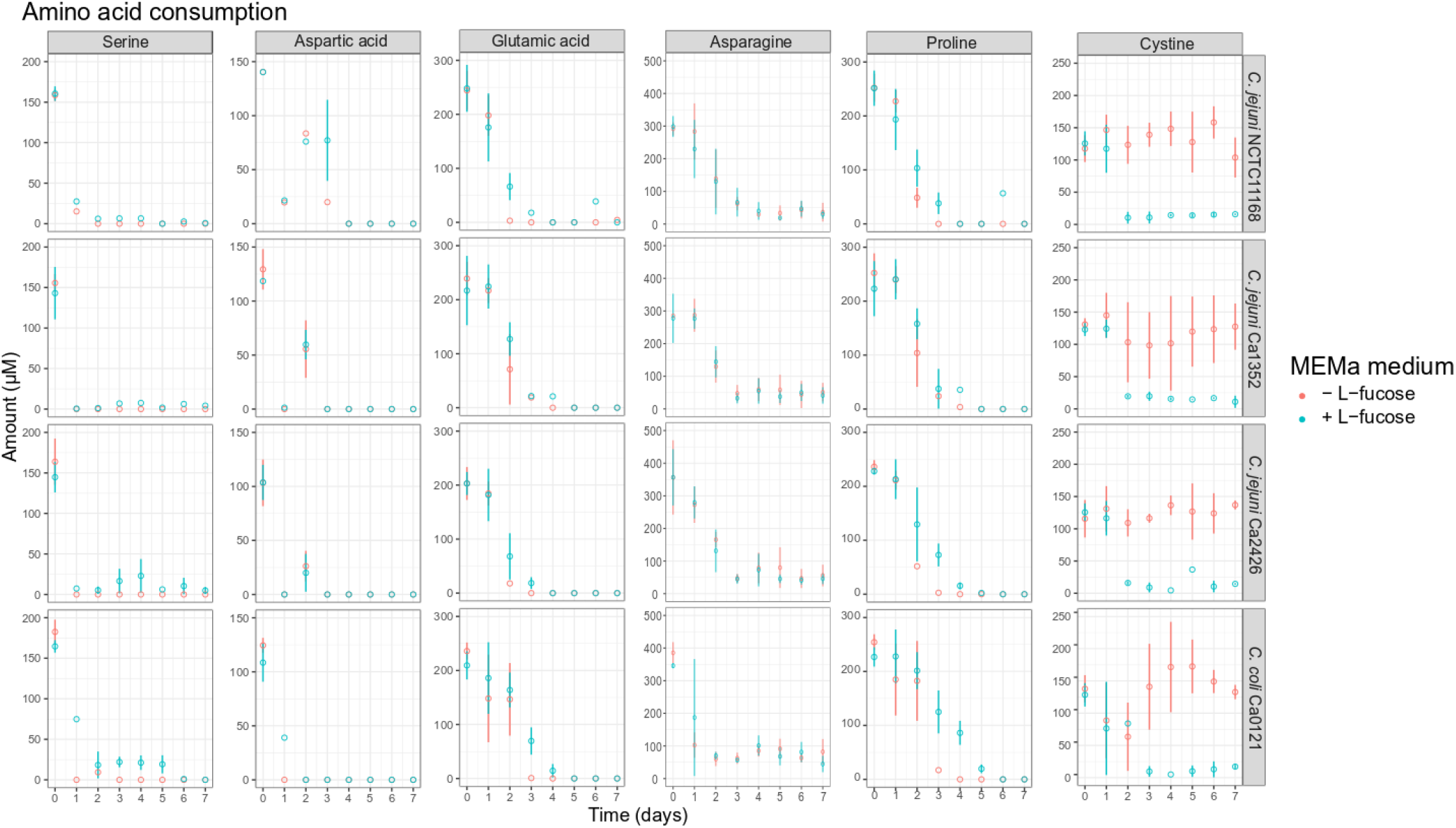
Amino acid production and consumption of selected *Campylobacter* isolates in MEMαF medium (blue dots) and MEMα medium (red dots) after 7-day incubation period. Columns correspond to different amino acids and each row shows results for a different isolate. Each value represents the average of three biologically independent replicates, and error bars show the standard deviation.

### Comparative analysis of L-fucose utilization clusters in *C. jejuni* and C. coli isolates

To investigate the genomic differences between the L-fucose utilization clusters of the tested isolates, we aligned each of the protein sequences (Cj0480c – Cj0490) of isolates *C. jejuni* Ca1352, Ca2426 and *C. coli* Ca0121 to the respective protein sequences of the reference isolate *C. jejuni* NCTC11168. The corresponding putative functions of the proteins encoded by the genes together constitute the L-fucose degradation pathway (Fig. 4A). Alignments of the gene *Cj0489*, encoding an aldehyde dehydrogenase, confirmed a previously discovered frameshift in the reference isolate NCTC11168 (36). However, this frameshift was not observed in *C. jejuni* Ca1352, Ca2426 and *C. coli* Ca0121 (Fig. 4B). The *Cj0489*-gene codes for a protein of 479 AA in *C. jejuni* Ca1352, Ca2426 and *C. coli* Ca0121, while the frameshift in the *C. jejuni* NCTC11168 genome, resulted in a shortened *Cj0489* gene, coding for a putative protein composed of 77 AA. Based on the presence of new start codon, a second larger fragment of the *Cj0489* gene, renamed *Cj0490* (36,37), is predicted to code for a putative protein of 394 AA. The AA composition of truncated proteins encoded by *Cj0489-Cj0490* of *C. jejuni* NCTC11168 was highly similar to the proteins encoded by the non-truncated *Cj0489* gene of *C. jejuni* Ca1352, Ca2426 and *C. coli* Ca0121, which was 95%, 95% and 97%, respectively. Next, we aligned all encoded proteins of the L-fucose cluster and calculated the percent identities to the reference isolate NCTC11168 (Fig. 4C and Table. S2). Only *Cj0482,* which encodes the altronate hydrolase/dehydratase, had 100% amino acid identity in all tested isolates. The altronate hydrolase/dehydratase encoded by *Cj0483* displayed >97% identity in all tested isolates. Clearly, the *C. jejuni* isolates Ca1352 and Ca2426 harbored more genes that displayed 100% protein identity with NCTC11168 than *C. coli* Ca0121. The L-fucose utilization cluster of *C. coli* Ca0121 was genetically the most distant from NCTC11168 with some translated genes having similarities as low as 84% such as the protein encoded by *Cj0487*.

**Figure 4.**
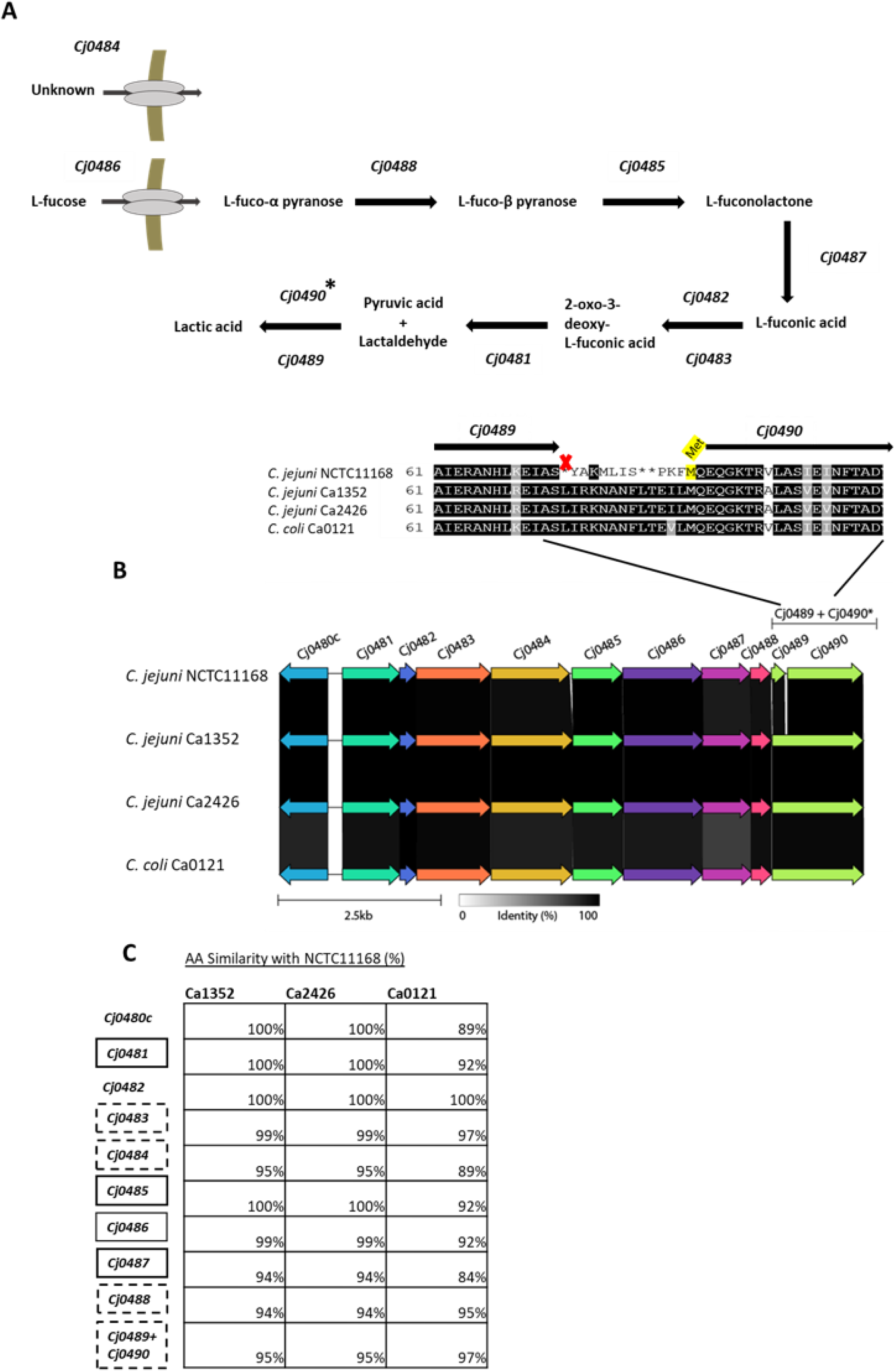
Genomic overview of the L-fucose utilization cluster in the four tested isolates. A) The predicted L-fucose metabolism cluster based on previously reported research (35,37). B) gene alignments performed with Clinker, AA identity is displayed in greyscale. A zoom-in is shown of the AA sequence of the *Cj0489-Cj0490* gene, indicating a frameshift in isolate NCTC11168, resulting in an early stop codon (marked with an X), splitting *Cj0489* (77 AA) and *Cj0490* (394 AA). A putative start site (Met) of C*j0490* in *C. jejuni* NCTC11168, is highlighted in yellow. C) Amino acid alignments of the genes in the isolates; *C. jejuni* Ca1352 (chicken meat isolate), *C. jejuni* Ca2426 (sheep manure isolate) and *C. coli* Ca0121 (pig manure isolate) with isolate *C. jejuni* NCTC11168 as reference (full alignment in Fig. S4). The percentages indicate AA similarities between *C. jejuni* NCTC11168 and Ca1352, Ca2426 or *C. coli* Ca0121. Similarity of *Cj0489-Cj0490* of isolate *C. jejuni* NCTC11168 is compared to *Cj0489* of the other isolates. Black and black dotted - boxes indicate genes to be essential or not for L-fucose metabolism based on mutant studies (34,35,37) (no data available for *Cj0482* and *Cj0480c*).

Notably, the proteins encoded by the L-fucose utilization cluster of *C. jejuni* Ca1352 and Ca2426 displayed 100% similarity with each other on amino acid level (Table. S2). Several SNPs were found between the 2 isolates, however, these did not result in single amino-acid polymorphisms.

## Discussion

The L-fucose utilization cluster is present in the majority of all *C. jejuni* and *C. coli* isolates, 65% and 73%, respectively, and is most commonly found in livestock-associated isolates (34). Comparative growth analysis of the four selected fuc+ *C. jejuni* and *C. coli* isolates showed that cell counts reached up to day 2 and 3 were similar in MEMα and MEMαF. This indicates, despite the initial onset of L-fucose consumption, that the base medium contains sufficient energy/carbon sources to support initial growth in the selected conditions. Impact of L-fucose on survival of the *C. jejuni* isolates, and the *C. coli* isolate, became apparent during prolonged incubation up to day 7, in line with the significant increase in L-fucose consumption in this period. Notably, only the *C. coli* isolate showed a further increase in cell counts from day 3 to day 7 in MEMαF.

In line with these observations, microscopy analysis of cells in non-supplemented MEMα medium showed in contrast to cells in MEMαF, an increased proportion of coccoid-shaped cells from day 3 to day 7. This points to a nutrient deficiency in this medium, in line with previous studies that showed starvation as a stress factor leading to a change from helical shaped to coccoid shaped cells (48–50). In addition, enhanced growth of the *C. coli* isolate was reflected in a more prominent fraction of spiral shaped cells morphology for a prolonged duration in MEMαF, pointing to a more robust phenotype for this isolate compared the tested *C. jejuni* isolates. Similar enhanced performance of the *C. coli* isolate was observed during growth in Mueller Hinton broths, MH2 and MH3 (see Fig. S2, containing 0.2% and 0.3% beef extract, respectively. No clear effect of the measured L-fucose consumption on growth performance in MH2 and MH3 was observed for the tested *C. jejuni* isolates, reaching maximum counts of ~8.0 log CFU/mL, while, *C. coli* Ca0121 reached highest CFU counts in both MH2 and MH3, and with added L-fucose, even ~9.5 log CFU/mL was reached in MH3 (Fig. S2).

HPLC analyses confirmed that the majority of the preferred amino acids were depleted on day 3 in MEMα and MEMαF, in line with previous studies that suggested that amino acids are preferred substrates for growth of *Campylobacter* (35). Notably, cystine, the oxidized dimer form of the amino acid cysteine, was only depleted at day 2 or day 3 when L-fucose was present in the medium and fucose consumption had started, indicating that cystine depletion is linked to L-fucose metabolism. In contrast, another study found production of sulfur-containing thiazolidines (methionine and cystine) upon addition of L-fucose to the medium, while no apparent link with L-fucose metabolism was found, led the authors to conclude that these sulfur-containing metabolites result from chemical reactions (38). The recent identification and characterization of a dedicated cystine transporter (*Cj0025c*) in *C. jejuni*, which is present in all our tested isolates (data not shown), next to the conceivable uptake of cystine via peptide transporters (30,51), may offer support for our observation that cystine depletion is linked to L-fucose metabolism in the tested *C. jejuni* and *C. coli* isolates in our study.

HPLC analyses of cultures grown in MEMα without and with added L-fucose demonstrated that acetate, lactate and pyruvate were mainly detected when *Campylobacter* was grown in MEMαF medium. Lactate and pyruvate can be metabolized by *Campylobacter* and are also hypothesized intermediates of the predicted L-fucose metabolism pathway, but their accumulation in the medium has not been shown before (35,52,53). Here we showed accumulation of pyruvate and lactate in the medium which is solely dependent on L-fucose, for isolates *C. jejuni* 1352 and 2426. For isolates *C. jejuni* NCTC11168 and *C. coli* Ca0121, on the other hand, pyruvate concentrations did not increase; they remained at low levels, in contrast with growth in MEMα medium where pyruvate is depleted after 1 day, suggesting that in isolates *C. jejuni* NCTC11168 and *C. coli* Ca0121 pyruvate is consumed at the same rate as it is produced. Interestingly, Van der Stel., et al. (2018) showed that catabolite repression in *C. jejuni* during growth on lactate and pyruvate, correlates with accumulation of intracellular succinate (54). In our study we observed a transient increase of succinate levels in the medium, with highest levels reached in MEMαF medium, in all tested isolates, conceivably linked to production and accumulating of lactate and pyruvate during L-fucose metabolism.

High concentrations of acetate were found over time in all tested isolates during growth in the presence of L-fucose. As acetate is a general by-product of *Campylobacter* metabolism, which is secreted when preferred carbon/energy sources are available (27), it is evident that L-fucose is preferred over acetate.

The L-fucose utilization cluster consists of 11 genes (*Cj0480c-Cj0490)* for *C. jejuni* NCTC 11168 and 10 genes (*Cj0480c-Cj0489)* for *C. jejuni* Ca1352, Ca2426 and *C. coli* Ca0121 and is predicted to metabolize L-fucose into lactic acid (35,37). Interestingly, the utilization cluster contains two sets of genes with putative overlapping functions, *Cj0482* and *Cj0483*, and *Cj0489* and *Cj0490*. Both *Cj0482* (88 AA) and *Cj0483* (389 AA), are two separated genes that are annotated as altronate hydrolases/dehydratases with similar ORFs in all tested isolates (Fig. 4). STRING analyses displayed a possible N-terminus (*Cj0482)* and possible C-terminus (*Cj0483)* of the predicted altronate hydrolase. In *C. jejuni*, only knockout studies of *Cj0483* have been performed, and this gene was found to be not essential for the metabolism of L-fucose (37). Stahl et al. (2011) also showed, by using microarrays, that both *Cj0482* and *Cj0483* were upregulated in the presence of L-fucose, 1.9 log_2_ and 4.4 log_2_ fold-change respectively (37). No knockout studies were performed with a *Cj0482* deletion mutant, however, due to the observed activity of this gene we hypothesize that both genes encode enzymes that can perform transformation of L-fuconate into 2-oxo-3-deoxy-L-fuconate. Comparative analysis shows that only in *C. jejuni* NCTC11168, the gene *Cj0489* presents a 1 bp deletion frameshift that results in an early stop, splitting *Cj0489* into *Cj0489* and *Cj0490*, both annotated as aldehyde dehydrogenase (36,55). In *C. jejuni* Ca1352, *C. jejuni* Ca2426 and *C. coli* Ca0121, *Cj0489* does not contain a frameshift and conceivably encodes the intact, 479 amino acid aldehyde dehydrogenase (Fig. 5B). Previous studies in *C. jejuni* NCTC11168 showed that deletion of *Cj0489* or *Cj0490* did not impair growth in the presence of L-fucose (34,35,37). However, in a transposon mutagenesis study using a hyper-invasive clinical isolate, *C. jejuni* isolate 01/51, three mutants were isolated that contained transposon insertions in the gene *Cj0490*, which showed impaired invasion of the human epithelial cell line INT-407 (55). Our HPLC data confirmed production of lactate in MEMαF, the final product of the L-fucose utilization pathway in all tested isolates, including NCTC11168, suggesting that L-fucose metabolism is not hampered in *C. jejuni* NCTC11168 that contains the frame shift. A recent study performed by Pascoe et al. (2019) reported the outcome of a domestication study analyzing 23 whole genome sequenced *C. jejuni* NCTC11168 isolates collected from a range of research laboratories across the UK (56). Our analysis of the L-fucose utilization cluster in these 23 *C. jejuni* NCTC11168 isolates showed that the fucose utilization clusters including the frameshift were 100% identical. This points to selection pressure on maintaining this L-fucose cluster with the frameshift in *Cj0489*. Whether the *Cj0489* and *Cj0490* genes in *C. jejuni* NCTC11168 encode (a) functional enzyme(s), or that there is an alternative lactaldehyde dehydrogenase induced in this isolate, remains to be elucidated.

Our comparative genotyping analysis of the four L-fucose utilization clusters, revealed that the cluster of *C. jejuni* NCTC11168 had several additional genomic differences in comparison with *C. jejuni* Ca1352, Ca2426 and *C. coli* Ca0121. Notably, genomic comparison analyses showed that the L-fucose utilization clusters of Ca1352 and Ca2426 were 100% similar, presenting only synonymous SNPs, in line with observed similarity in phenotypic behavior of these two isolates. The L-fucose utilization cluster of *C. coli* Ca0121 was the most distant from the cluster of *C. jejuni* NCTC11168. Our results showed that, in the presence of L-fucose, *C. coli* Ca0121 was able to reach the highest CFU counts, maintained a spiral morphology and completely metabolized available L-fucose in the tested conditions. These results suggest a possible link between the observed *C. coli* Ca0121 phenotype and changes in amino acid composition of enzymes in the L-fucose utilization cluster that could point to altered enzyme levels and/or functionality. However, it should be noted that the housekeeping genes of *C. jejuni* and *C. coli* share 86.5% nucleotide sequence identity and that differences in growth, morphology, survival and metabolism may also be influenced by other genomic differences (57,58). Future studies should also include the role of L-fucose utilization pathway in the metabolism of D-arabinose (35). As D-arabinose is a rare sugar in nature and a high preference to L-fucose over D-arabinose was observed (35), it remains to be validated that D-arabinose is a natural substrate for the L-fucose utilization cluster.

In conclusion, our study demonstrated that possessing the L-fucose cluster is not only beneficial to *C. jejuni* NCTC11168 but also to other tested *C. jejuni* isolates Ca1352 and Ca2426, and *C. coli* isolate Ca0121. All tested isolates isolated from different hosts, showed enhanced survival and prolonged spiral shaped morphology in the presence of L-fucose, with the *C. coli* isolate having the most robust phenotype. Several genetic differences were observed between the gene clusters, however, it is uncertain whether these differences are linked to differences observed in phenotypic behavior. Further research into *Campylobacter* species and isolate diversity may reveal whether specific L-fucose utilization cluster genotypes link with specific phenotypic behavior including environmental transmission and between host species, and pathogenesis.

## Acknowledgements

This research was financially supported by the Graduate School VLAG (Wageningen University & Research, Wageningen, The Netherlands).

AZ acknowledges financial support by the Netherlands’ Organization for Health Research and Development (ZonMw) with grant number 50-52200-98-316 (project name: “DEPiCT – Discerning Environmental Pathways of Campylobacter Transmission”).

*C. jejuni* Ca1352 isolated from chicken meat, *C. jejuni* Ca2426 isolated from sheep manure and *C. jejuni* Ca0121 isolated from pig manure were obtained from Netherlands Food and Consumer Product Safety Authority.

## Supplementary

**Table. S1.** Components of 41061 - MEM alpha, nucleosides, no phenol red.

| Components | mM | Components | mM |
| --- | --- | --- | --- |
| **<u>Amino Acids</u>** |  | **<u>Inorganic Salts</u>** |  |
|  |  | Calcium Chloride (CaCl<sub>2</sub>) |  |
| Glycine | 0.67 | (anhyd.) | 1.80 |
|  |  | Magnesium Sulfate (MgSO<sub>4</sub>) |  |
| L-Alanine | 0.28 | (anhyd.) | 0.81 |
| L-Arginine | 0.49 | Potassium Chloride (KCl) | 5.33 |
|  |  | Sodium Bicarbonate |  |
| L-Asparagine-H<sub>2</sub>O | 0.33 | (NaHCO<sub>3</sub>) | 26.19 |
| L-Aspartic acid | 0.23 | Sodium Chloride (NaCl) | 117.24 |
| L-Cysteine |  | Sodium Phosphate monobasic |  |
| hydrochloride-H<sub>2</sub>O | 0.57 | (NaH<sub>2</sub>PO<sub>4</sub>-H<sub>2</sub>O) | 1.01 |
| L-Cystine 2HCl | 0.10 | **<u>Ribonucleosides</u>** |  |
| L-Glutamic Acid | 0.51 | Adenosine | 0.037 |
| L-Glutamine | 2.0 | Cytidine | 0.041 |
| L-Histidine | 0.2 | Guanosine | 0.035 |
| L-Isoleucine | 0.4 | Uridine | 0.040 |
| L-Leucine | 0.40 | **<u>Deoxyribonucleosides</u>** |  |
| L-Lysine | 0.40 | 2'Deoxyadenosine | 0.039 |
| L-Methionine | 0.10 | 2'Deoxycytidine HCl | 0.042 |
| L-Phenylalanine | 0.19 | 2'Deoxyguanosine | 0.037 |
| L-Proline | 0.35 | Thymidine | 0.041 |
| L-Serine | 0.24 | **<u>Other Components</u>** |  |
| L-Threonine | 0.40 | D-Glucose (Dextrose) | 5.55 |
| L-Tryptophan | 0.05 | Lipoic Acid | 9.71E-04 |
| L-Tyrosine |  |  |  |
| disodium salt | 0.23 | Sodium Pyruvate | 1.00 |
| L-Valine | 0.39 |  |  |
| **<u>Vitamins</u>** |  |  |  |
| Ascorbic Acid | 0.28 |  |  |
| Biotin | 4.10E-04 |  |  |
| Choline chloride | 0.01 |  |  |
| D-Calcium |  |  |  |
| pantothenate | 0.002 |  |  |
| Folic Acid | 0.002 |  |  |
| Niacinamide | 0.008 |  |  |
| Pyridoxal |  |  |  |
| hydrochloride | 0.005 |  |  |
| Riboflavin | 2.66E-04 |  |  |
| Thiamine |  |  |  |
| hydrochloride | 0.003 |  |  |
| Vitamin B12 | 0.001 |  |  |
| i-Inositol | 0.01 |  |  |

**Table. S2.** Overview of genes in the L-fucose utilization cluster.

| Essential genes | Function | Length in AA | AA Similarity with NCTC11168 (%) |  |  |
| --- | --- | --- | --- | --- | --- |
|  |  | NCTC11168/Ca1352/Ca2426/Ca0121 | Ca1352 | Ca2426 | Ca0121 |
| Cj0486 | Transporter | 418/418/418/415 | 99% | 99% | 92% |
| Cj0485 | Dehydrogenase | 263/263/263/263 | 100% | 100% | 92% |
| Cj0487 | Hydrolase | 256/256/256/256 | 94% | 94% | 84% |
| Cj0481 | DHDPS | 301/301/301/301 | 100% | 100% | 92% |
| **Non essential genes** |  |  |  |  |  |
| Cj488 | Mutarotase | 105/105/105/105 | 94% | 94% | 95% |
| Cj0482 | Altronate hydrolase/dehydratase | 88/88/88/88 | 100% | 100% | 100% |
| Cj0483 | Altronate hydrolase/dehydratase | 389/389/389/389 | 99% | 99% | 97% |
| Cj0489 | Dehydrogenase G | 77/479/479/479 | 14% | 14% | 15% |
| Cj0490 | Dehydrogenase | 129/129/129/129 | 100% | 100% | 99% |
| **Extra genes in cluster** |  |  |  |  |  |
| Cj0484 | Transporter | 411/429/429/424 | 95% | 95% | 89% |
| Cj0480c | L-fucose operon regulator | 253/253/253/253 | 100% | 100% | 89% |

**Fig. S1.**
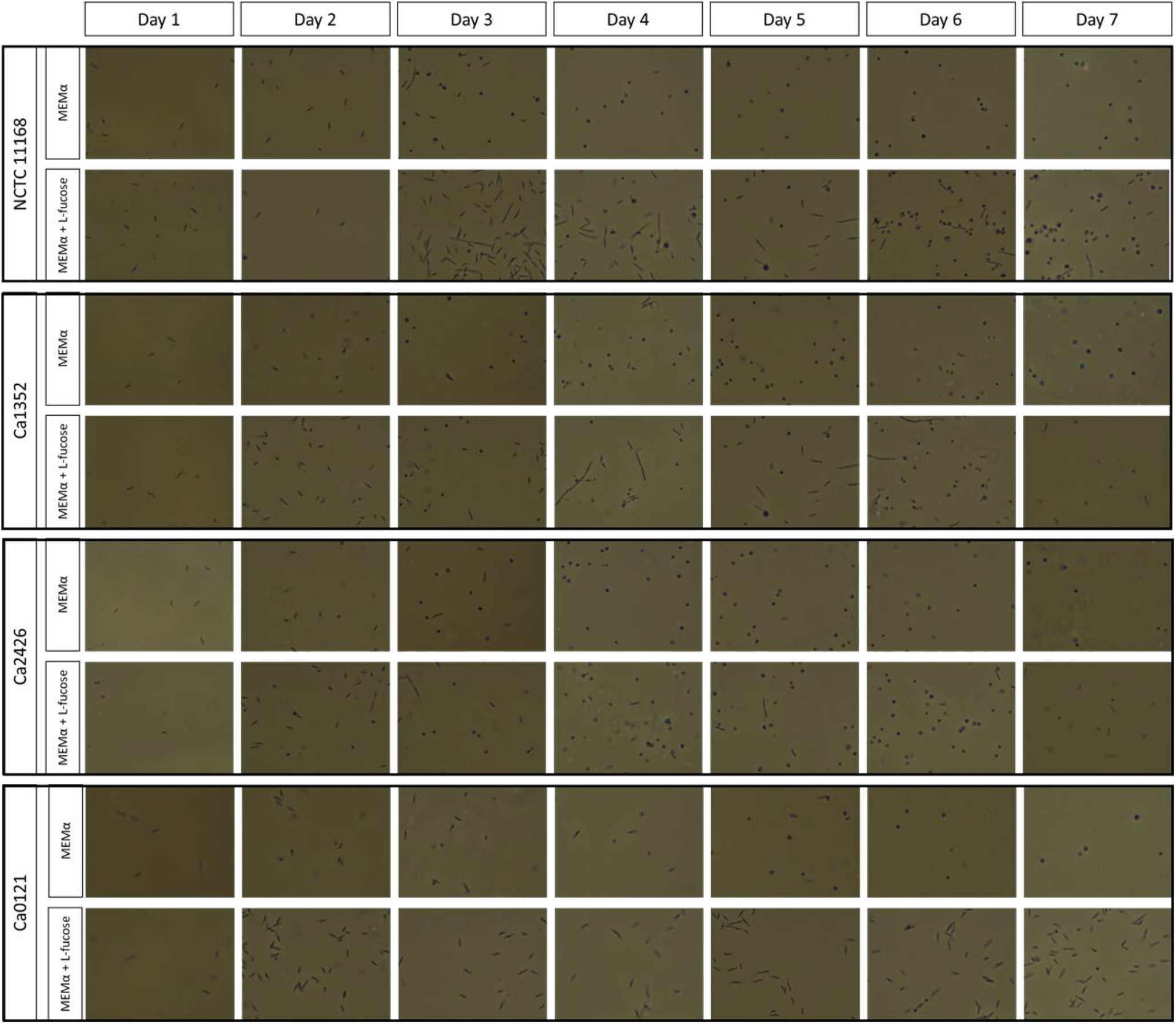
Microscopic images of the tested *Campylobacter* isolates. *Figure S1* Morphology of *C. jejuni* NCTC11168, Ca1352, Ca2426 and *C. coli* Ca0121 during a 7 day growth experiment in MEMα or MEMαF medium.

**Fig. S2.**
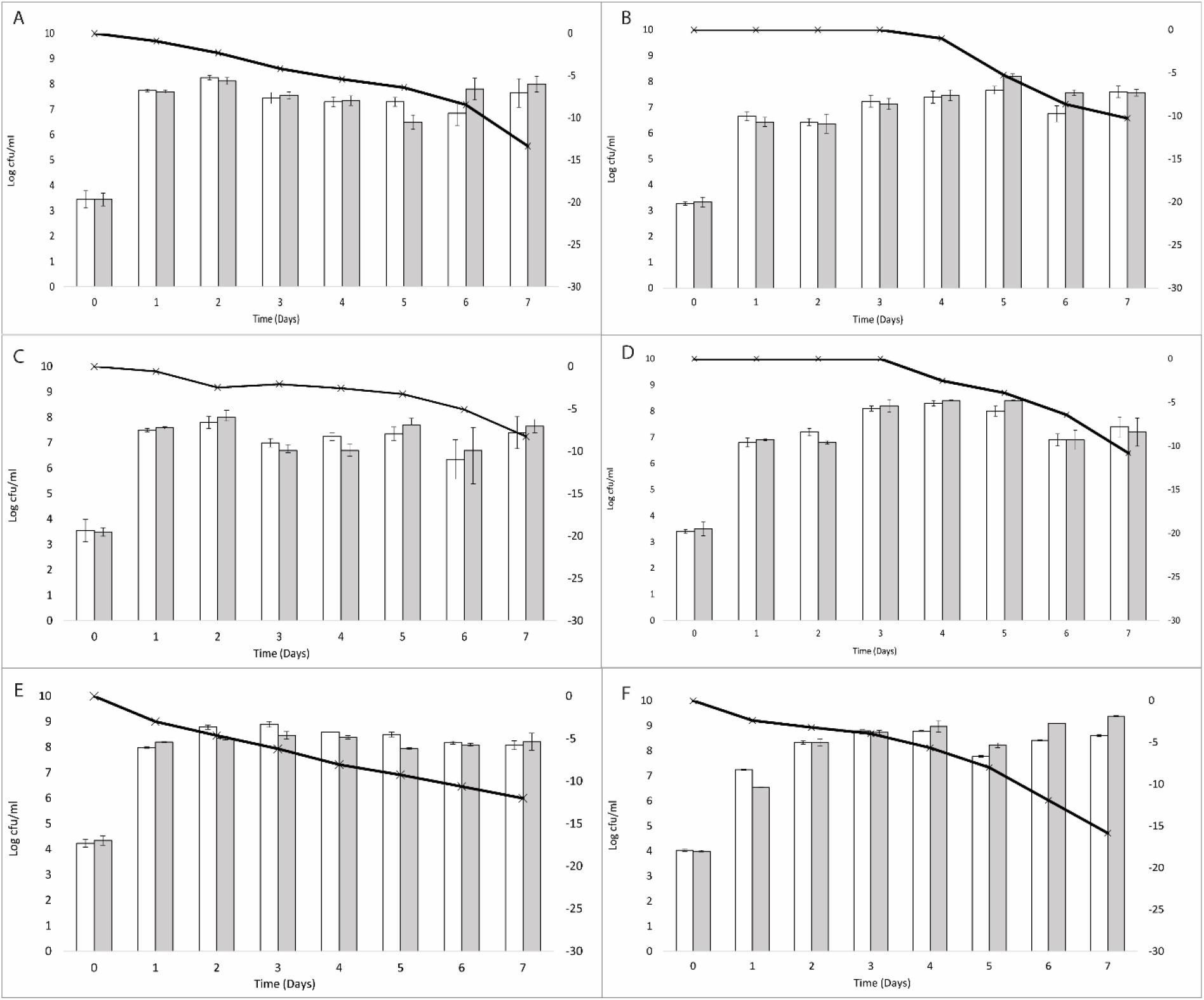
Growth and L-fucose consumption of *C. jejuni* NCTC11168, Ca2426 and *C. coli* Ca0121 in MH2 and MH3 medium. *Figure S2.* Quantification of planktonic growth for **A)** isolate *C. jejuni* NCTC11168 in MH2 medium, **B)** *C. jejuni* NCTC11168 in MH3 medium **C)** *C. jejuni* Ca2426 in MH2 medium, **D)** *C. jejuni* Ca2426 in MH2 medium, **E)** *C. coli* Ca0121 in MH2 medium and **F)** *C. coli* Ca0121 in MH3 medium. White bars represent MH2 or MH3 medium and grey bars represent MH2 or MH3 medium + L-fucose. The black line shows consumption of L-fucose over time.

**Fig. S3.**
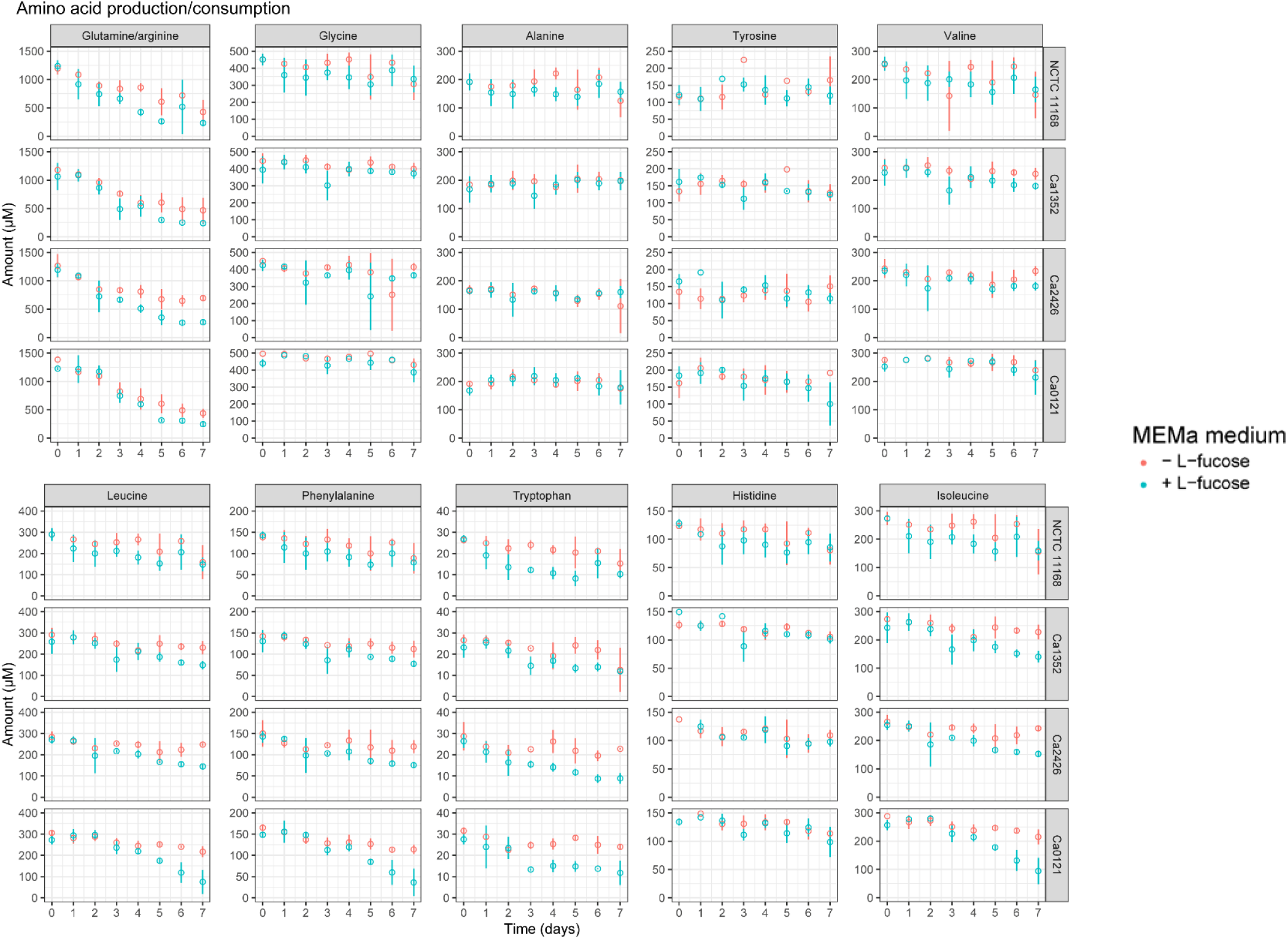
HPLC measurements of amino acids. *Figure 2* Amino acid production and consumption of selected *Campylobacter* isolates in MEMαF medium (blue dots) and MEMα medium (red dots) after 7-day incubation period. Columns correspond to different amino acids and each row shows results for a different isolate. Each value represents the average of three biologically independent replicates, and error bars show the standard deviation.

**Fig. S4.**
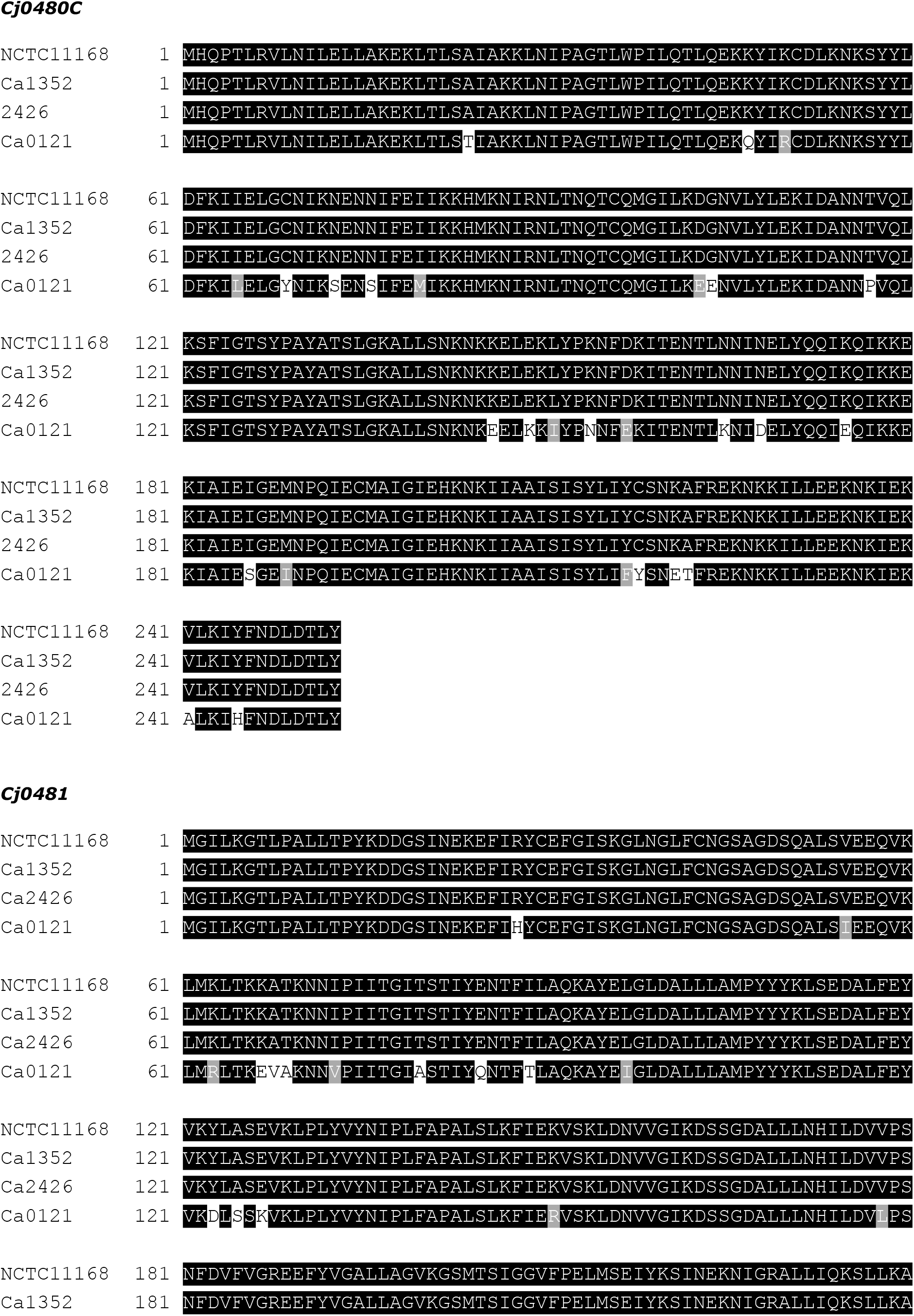
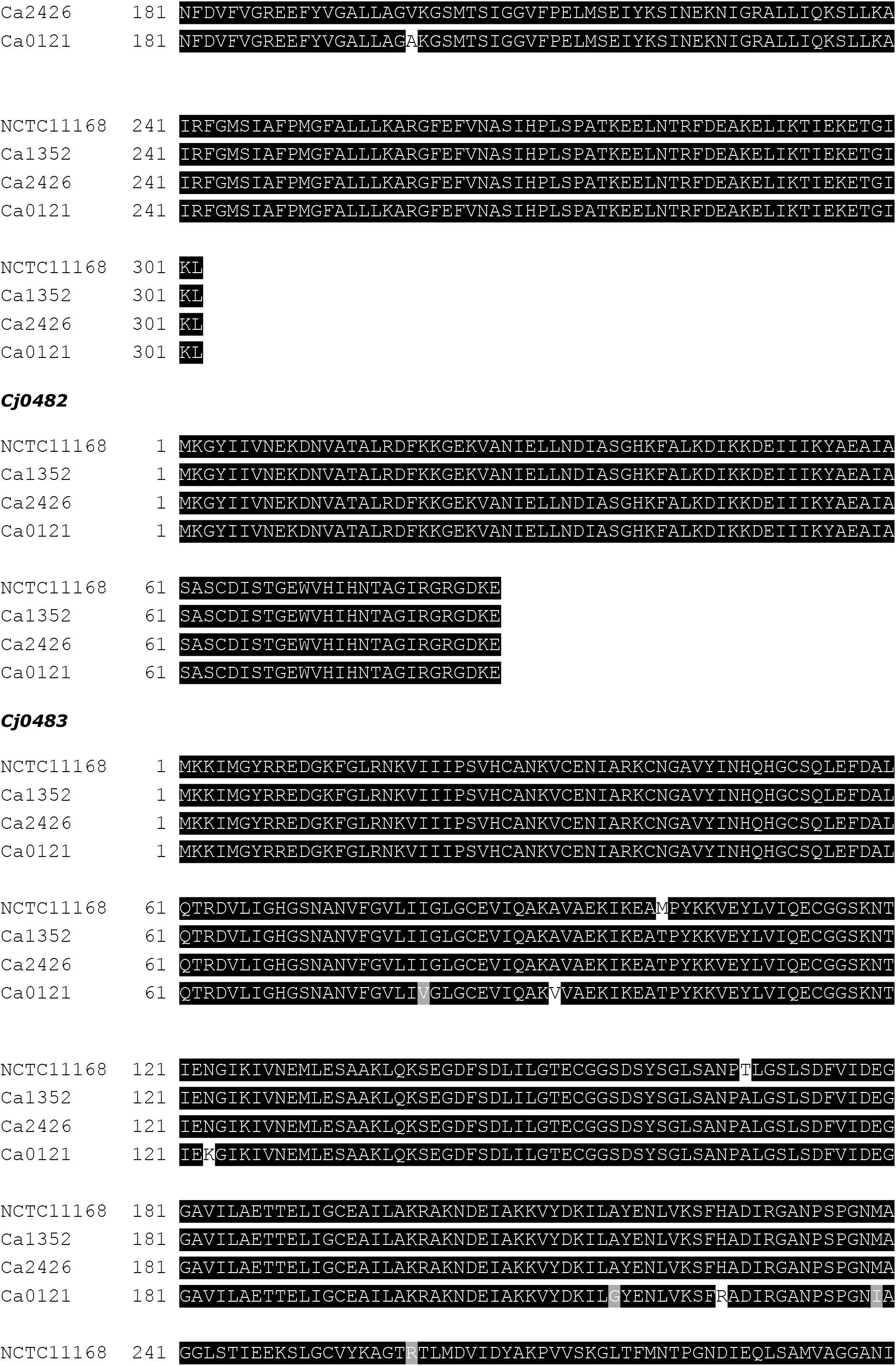
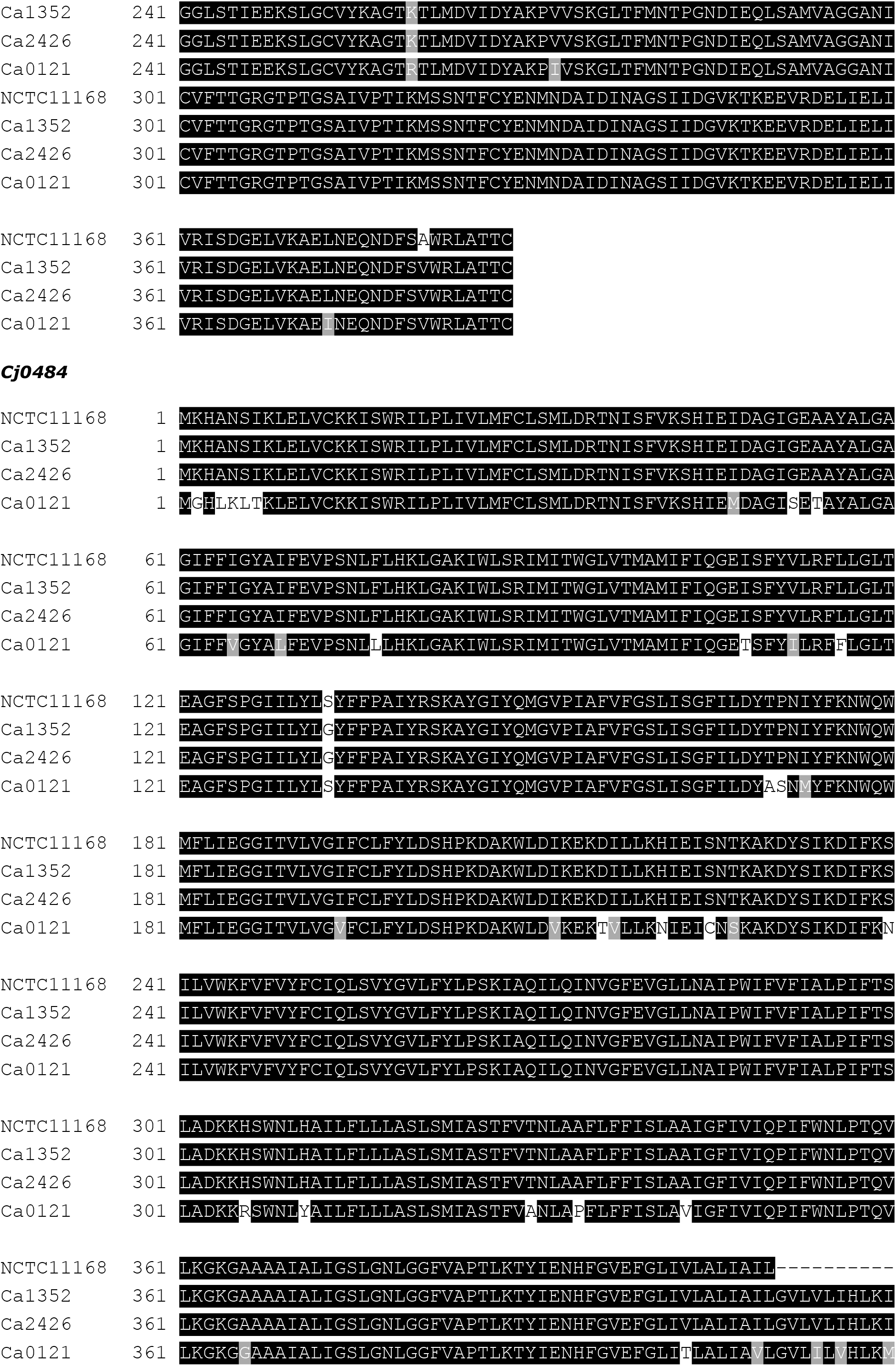
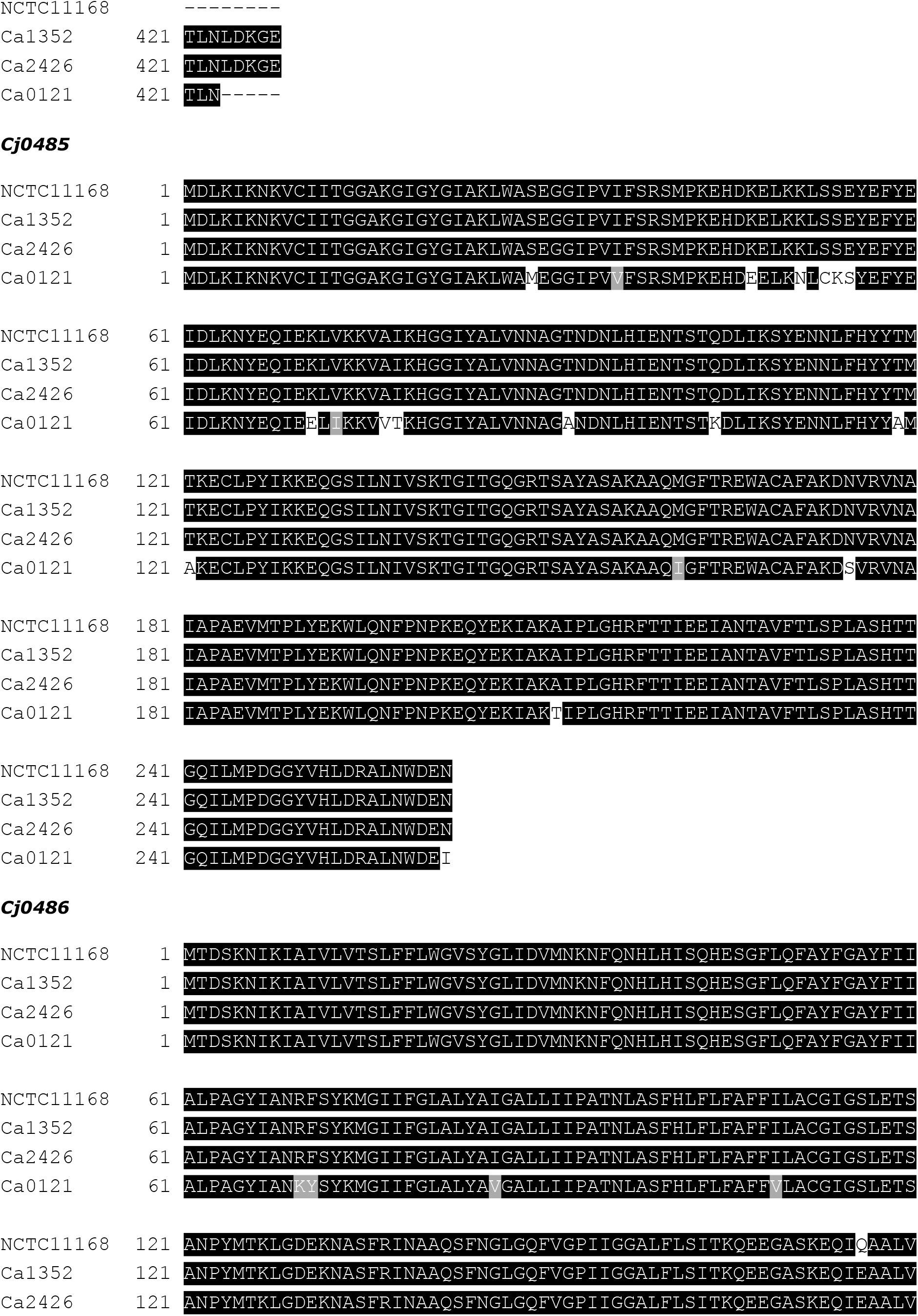
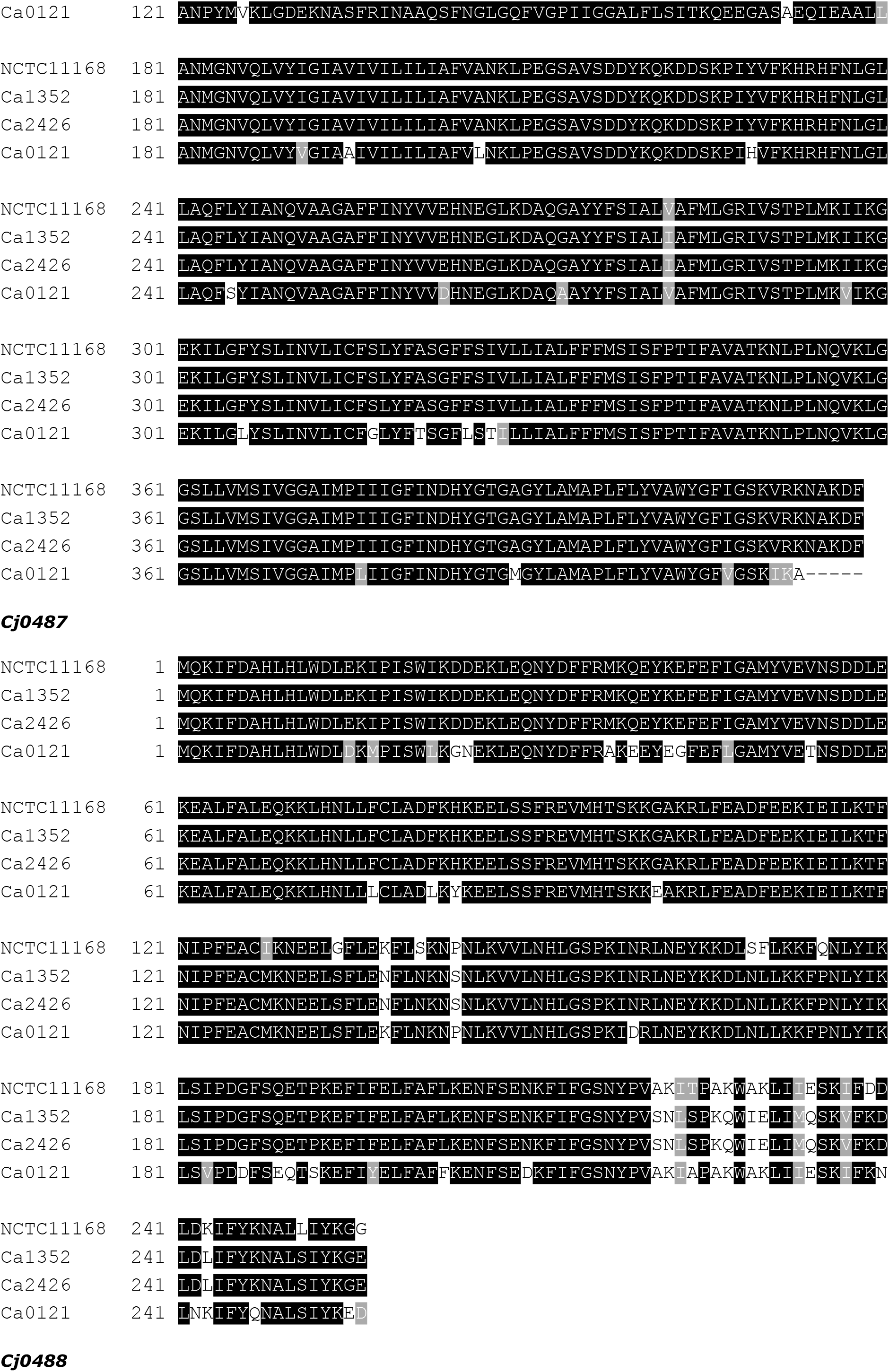
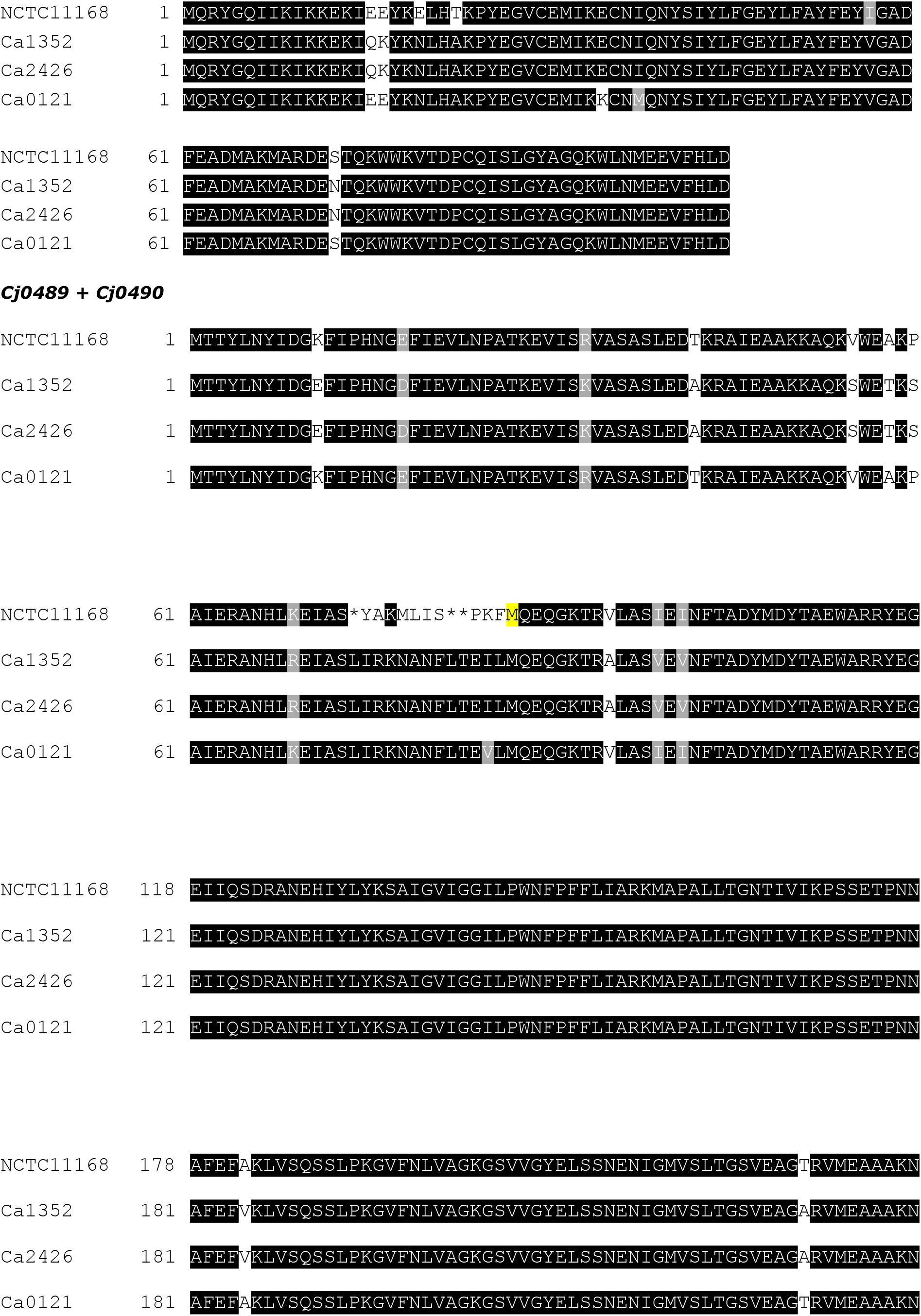
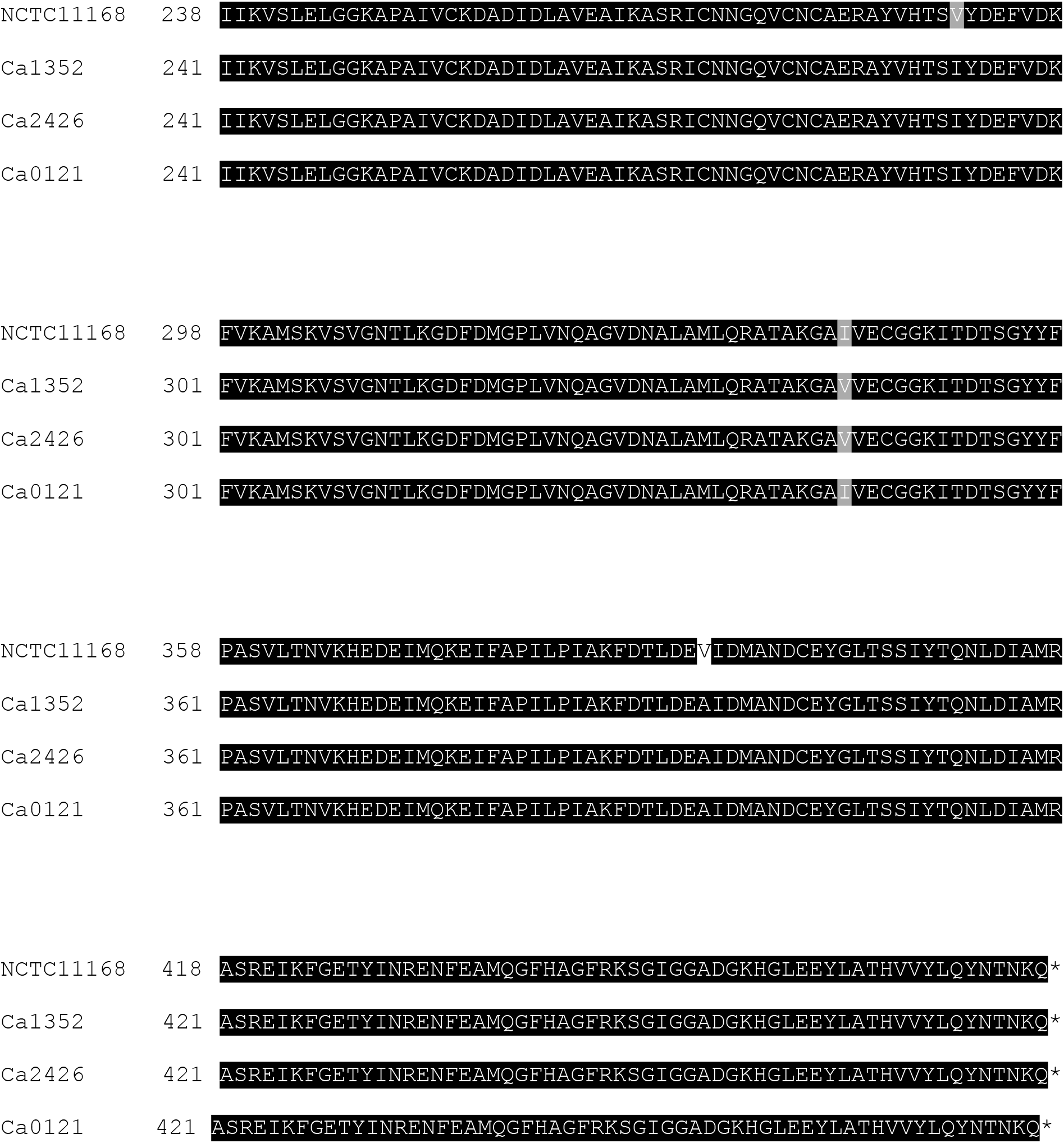
Complete L-fucose gene cluster alignment.

